# Overground gait adaptability in older adults with diabetes in response to virtual targets and physical obstacles

**DOI:** 10.1101/2022.10.19.512897

**Authors:** Suzanne Martin, Simon B. Taylor, Blynn L. Shideler, Rajna Ogrin, Rezaul Begg

**Author notes:** Suzanne Martin, PhD, Research Fellow (Corresponding Author), Institute of Health Exercise and Sport, Victoria University, Ballarat Rd, Footscray, Victoria, Australia.

## Abstract

**Background:** To step over an unexpected obstacle, individuals adapt gait; they adjust step length in the anterior-posterior direction prior to the obstacle and minimum toe clearance height in the vertical direction. Inability to adapt gait may lead to falls in older adults with diabetes. Therefore, this study aimed to investigate gait adaptability in older adults with diabetes.

**Research question:** Does diabetes impair gait adaptability and increase sagittal foot adjustment errors?

**Methods:** Three cohorts of 16 people were recruited: young adults (Group I), healthy older adults (Group II), and older adults with diabetes (Group III). Participants walked in baseline at their comfortable speeds. They then walked and responded to what was presented in gait adaptability tests which included 40 trials with four random conditions: step shortening, step lengthening, obstacle avoiding, and walking through. Virtual step length targets were 40% of the baseline step length longer or shorter than the mean baseline step length; the actual obstacle was a 5-cm height across the walkway. A Vicon three-dimensional motion capture system and four A.M.T.I force plates were used to quantify spatiotemporal parameters of a gait cycle and sagittal foot adjustment errors (differences between desired and actual responses in the second step of the gait cycle). Analyses of variance (ANOVA) repeated measured tests were used to investigate group and condition effects on dependent gait parameters at a significance level of 0.05.

**Results:** Statistical analyses of Group I (n = 16), Group II (n = 14) and Group III (n = 13) revealed that gait parameters did not differ between groups in baseline. However, they were significantly different in adaptability tests. Group III significantly increased their stance and double support times in adaptability tests, but these adaptations did not improve their foot adjustments. They had the greatest step length errors and the lowest toe-obstacle clearance which might cause them to touch the obstacle the most.

**Significance:** The presented gait adaptability tests may serve as entry tests for falls prevention programs.

## Introduction

In older populations (aged 65 or older), there is an increased risk of falls in individuals with diabetes [1]. The annual incidence of falls has been reported to be 39% in older people with diabetes [2]. Meanwhile, over 50% of older people with diabetes report at least one injurious fall or two non-injurious falls a year [3]. Around 30-50% of these falls caused minor injuries, and 5-10% caused major injuries including fractured neck of femur [4].

Apart from fall-related injuries, falls greatly increase the cost of national health care. The reported direct annual medical cost was US$23.3 billion in the United States and US$1.6 billion in the United Kingdom [5]. Fall-related events account for 40% of nursing home placements, and contribute to further increases in healthcare costs [6].

Impaired gait adaptability is associated in falls in some pathological populations [7]. Gait adaptability refers to altering gait according to changes in the environment to adjust the displacement of the foot in response to goal-oriented tasks [8–10]. Gait adaptability impairs in healthy older adults while responding to goal-oriented tasks [8, 11–14]. Impaired gait adaptability causes a contact between the foot and obstacles during obstacle avoidance [15]. However, people with impaired gait adaptability are able to avoid obstacles if they see the obstacles in advance so they will have enough time to adjust their gait when responding. The adaptation of step lengths and foot-obstacle clearance are apparent results of early gait adaptability in previous studies [8, 12, 16, 17] in which participants adjusted their step length a few steps prior responding to a stepping target or an obstacle.

Despite the prevalence of falls in older adults with diabetes, only a few studies [18–20] have investigated the effects of diabetes in response to obstacle avoidance. Although these studies reported that diabetes reduced toe-obstacle clearances, none of their participants touched obstacles [18, 19]. This implies that when obstacles are always visible and participants can see obstacles from the starting point, they can adapt the lengths of steps a few steps ahead to place obstacles in the middle of their crossing steps and avoid them. In dynamic environment, there is a time when a person needs to respond suddenly so the present study aimed to investigate responses of older people with diabetes to tasks which suddenly and randomly appeared. To do so, three groups of young adults, healthy older adults and older adults with type two diabetes (T2D) were tested to investigate gait with and without challenges. It was hypothesised that diabetes, ageing, and the combination of diabetes and ageing would impair gait with and without challenges.

## Methods

### Participants

A priori power calculation estimated 48 participants in three group would be required to detect an effect size of 0.66 with 95% power. Therefore, 16 young adults, 16 healthy older adults, and 16 older adults who lived with T2D for at least five years without having diabetic complications (i.e. foot ulcer) were recruited. All participants had no cognitive function impairment (The mini-mental state examination (MMSE) [21] score ≥ 23), vision impairment, and ageing- and diabetes-related neuropathy (The Michigan neuropathy screening instrument [22] score ≤ 3).

All procedures were approved by the Victoria University Human Research Ethics Committee (HRE17-194). All participants provided written informed consent; data was collected at the Institute for Health and Sport, Victoria University, Melbourne, Australia.

### Experimental set-up and procedure

A three-dimensional motion analysis system (VICON, Oxford, UK) with 14 cameras and four A.M.T.I forces plates (Advanced Mechanical Technology Inc., Watertown, Mass., USA) was used to record gait biomechanics. The motion analysis system tracked four total reflective markers, attached bilaterally to athletic shoes over the first metatarsals and heels of each foot. The system recorded the ground reaction forces when participants walked for eight metres at their self-selected speed in baseline without challenge and in adaptability tests with four challenging conditions.

Participants walked in a baseline walking condition (6 trials) and then completed adaptability tests (40 trials). Adaptability tests entailed participants manoeuvring one of four challenging conditions: shortening the target step, lengthening the target step, crossing an obstacle, or walking through without any change. Participants were blinded to the gait adaptability challenge during any given trial; challenges were randomly assigned and presented to participants two steps ahead during overground walking (Figure 1).

**Figure 1.**
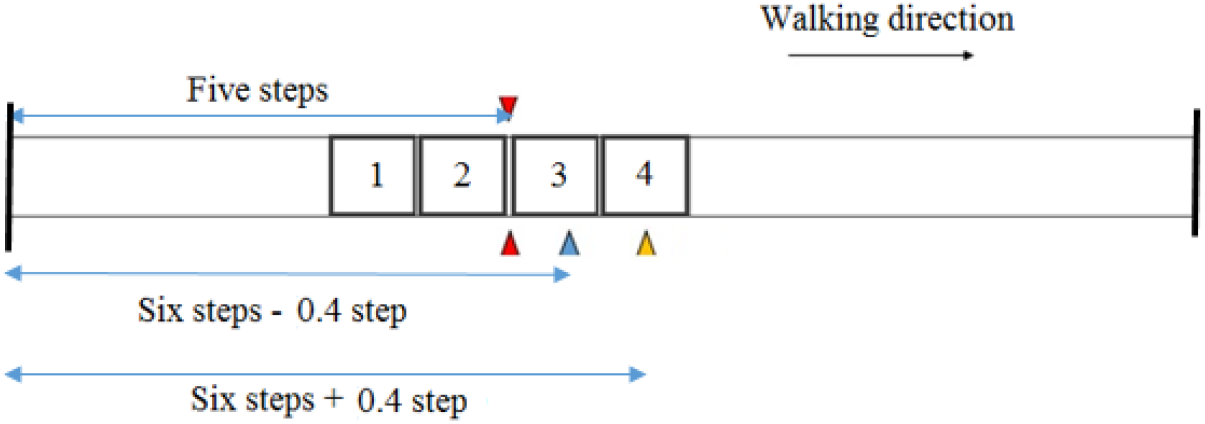
Overhead view of the experimental setup in the foot displacement adaptation test in the walkthrough condition. Red triangles present locations of the potential obstacle. The blue and yellow triangles present the locations of two laser beam projectors for potential short and long stepping targets, respectively. Two black lines present the starting and finishing points.

Short and long step targets (SSL and LSL targets) were projected on the walkway by using laser beams virtually whereas the crossing condition was presented two steps ahead of a participant by a fine cord located on the ground which was lifted 5 cm by two servomotors (Figure 2).

**Figure 2.**
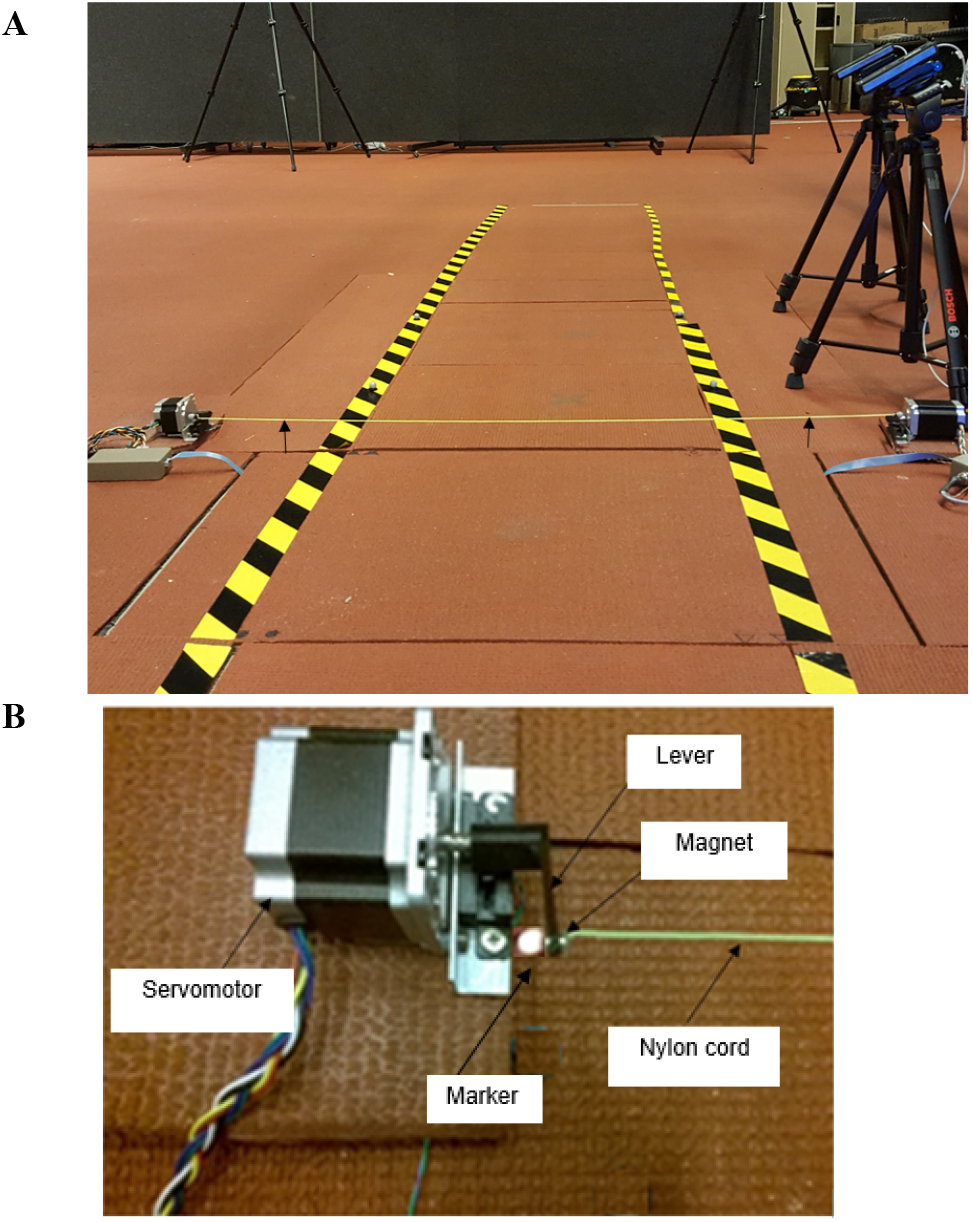
The 5-cm low height obstacle was presented sometimes in gait adaptability tests. The yellow nylon cord was held at a 5-cm height when the synchronised servomotors were turned on by touching the first force platform (A). A nylon cord was attached on each end to a servomotor’s lever by a small magnet and sat on the ground when participants stood behind the starting points (five steps away) and was randomly lifted in the air to show a 5-cm height when participants were two steps away (B).

Step targets were scaled to a participant’s baseline step length (SL) and were 0.6 × baseline SL and 1.4 × baseline SL [23].

A slight touch of a foot during crossing the obstacle immediately caused the cord attached to the servomotors to become detached. Thus, the forward swing of the leg was not affected balance if the participants could not avoid the obstacle.

A condition in which neither a short/long step target nor the obstacle was presented, was also added to investigate the effect of knowing something might be presented on gait.

Experimental trials were deemed successful if participants took five steps, responded to a condition in their sixth steps, and continued walking until they passed the finishing point.

### Data processing and analysis

Nexus Vicon software was used to analyse individual gait cycles (defined as the heel strike of a foot to the heel strike of the same foot) in each trial in the baseline and adaptability tests. Spatiotemporal parameters (step velocity, stance time, swing time, double support (DS) time, step length (SL)) were computed. Leg length for each participant was used to normalise the previous and target step length (the first and second step of the gait cycle). Velocity (m/s) was calculated by dividing SL to the step time (SL/step time). Stance time, swing time and DS time were reported as percentages of a gait cycle. The horizontal distance between the toe marker and projected laser lines in the target step were called long/short step length absolute errors (errors) and the vertical distance between the toe marker and the obstacle in the target step was called toe-obstacle clearance heights in adaptability tests.

### Statistical analysis

All statistical analyses (α = 0.05) were performed in SPSS (Version 25 for Windows, SPSS Science, Chicago, USA). Analyses of variance (ANOVA) repeated measured tests were used to investigate the effects of group (young vs. healthy older vs. older diabetes) and condition (baseline vs. walkthrough vs. short/long/obstacle) on dependent gait parameters (velocity, stance time, swing time, double support time, step length). Multiple comparisons (Bonferroni post hoc tests) were used when the main effect of groups was significant. Nonparametric tests (Kruskal-Wallis H and Mann-Whitney U) were used to compare short/long step length errors and toe obstacle clear heights between groups.

## Results

Forty-three participants (16 persons in Group I, 14 persons in group II, 13 persons in Group III) completed gait adaptability tests. Excluded participants included two participants of Group II and one participant of Group III who felt uncomfortable to walk on the treadmill and two participants of Group III who had a history of fall.

Table 1 presents the characteristics of participants. Older participants in Group II and Group III were significantly older and taller than young participants; however, no significant differences were found between characteristics of Group II and Group III. Group I differed from Group II and Group III in terms of age (*p* < 0.001 and *p* < 0.001) and height (*p* = 0.034 and *p* = 0.039).

**Table 1.**
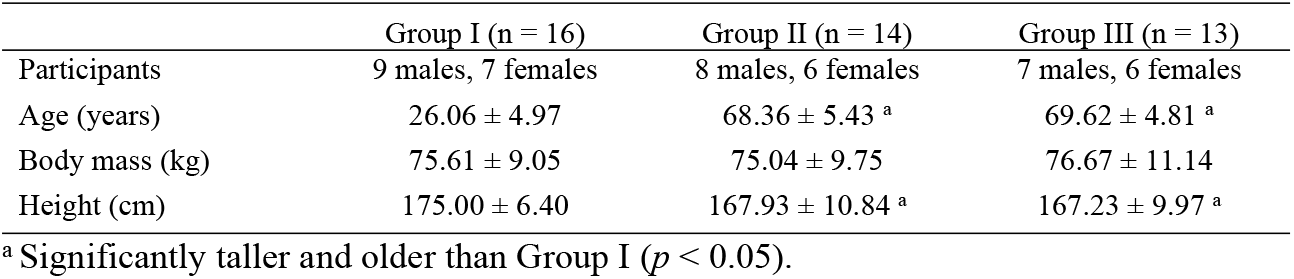
Descriptive statistics of each group pf participants. Mean ± standard deviation of age, body mass and height in young adults (Group I), healthy older adults (Group II), and older adults with diabetes (Group III) are presented.

Table 2 presents target step length absolute errors and toe-obstacle clearance heights of participants during adaptability tests. Mean absolute errors in response to SSL and LSL targets were different between groups (H (2) = 25.05, *p* < 0.0001, and H (2) = 24.13, *p* < 0.0001). Absolute errors were different between Group I and Group III (U = 3, p < 0.0001 and U = 5, p < 0.0001) and Group II and Group III (U = 7.5, p < 0.0001 and U = 6, p < 0.0001). Between groups, only toe-obstacle clearance heights were significantly different (H (2) = 10.03, *p* = 0.007). The mean toe-obstacle clearance height of Group III was significantly shorter than those of both Group I (U = 44, *p* = 0.007) and Group II (U = 33, *p* = 0.004).

**Table 2.** Sagittal foot displacement adaptation absolute errors in three groups of participants. Group I, young adults (n =16); Group II, healthy older adults (n = 14); and Group III, older adults with diabetes (n = 13). Mean ± standard deviation of absolute errors are presented.

|  | Step length error (cm) |  | Toe-obstacle vertical distance (cm) |
| --- | --- | --- | --- |
|  | Shortening | Lengthening |  |
| Group I | 2.34 ± 0.45 | 1.58 ± 0.56 | 7.95 ± 2.86 |
| Group II | 2.58 ± 0.52 | 1.75 ± 0.54 | 7.79 ± 2.01 |
| Group III | 4.69 ± 1.41 <sup>a,b</sup> | 3.95 ± 1.44 <sup>a,b</sup> | 5.46 ± 3.39 <sup>a,b</sup> |

<sup>a</sup> Significantly different from the healthy older group ( $p < 0.05$ ).
<sup>b</sup> Significantly different from the young group ( $p < 0.05$ ).

Table 3 presents gait parameters in baseline and in adaptability tests in the challenging conditions.

**Table 3.**
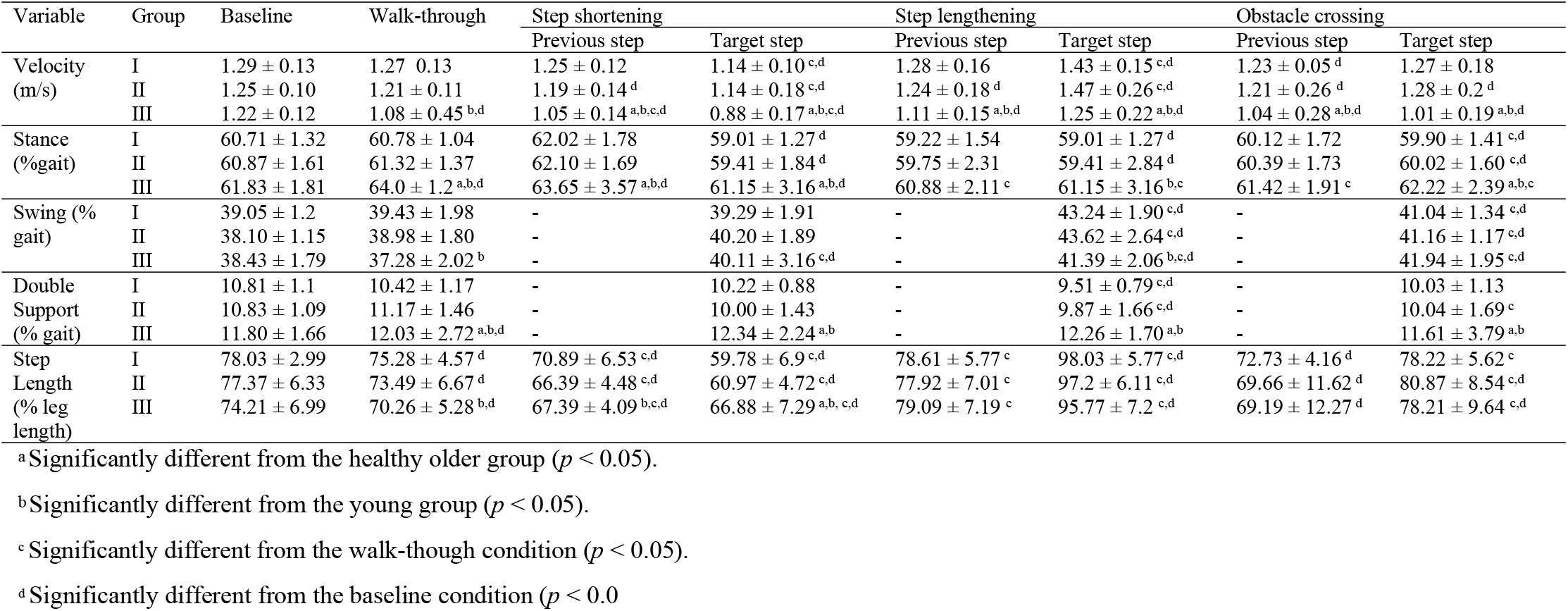
The comparison of gait and gait adaptability between groups. Mean ± standard deviation (SD) of spatiotemporal parameters in Group I, young adults (n =16); Group II, healthy older adults (n = 14); and Group III, older adults with diabetes (n = 13) in baseline walking and adaptability test conditions.

### Step shortening condition

Conditions were significantly different in target step velocity (*F*_1.65, 65.98_ = 50.66, *p* < 0.001); previous step velocity (*F*_1.65, 66.33_ = 19.17, *p* < 0.001); leading leg stance time (*F*_2,80_ = 14.51, *p* < 0.001); swing time (*F*_2,80_ = 6.422, *p* = 0.003); DS time (*F*_2,80_ = 6.97, *p* = 0.002); target step length (*F*_1.69, 67.85_= 109.68,*p* < 0.001); and previous step length (*F*_2,80_ = 63.76,*p* < 0.001).

Groups were significantly different in target step velocity (*F*_2, 40_ = 10.64, *p* < 0.001), previous step velocity (*F*_2,40_ = 7.11, *p* = 0.002), trailing leg stance time (*F*_2,40_ = 6.85, *p* = 0.003), leading leg stance time (*F*_2,40_ = 6.95,*p* = 0.003), DS time (*F*_2,40_ = 9.29,*p* < 0.001,), previous step length (*F*_2,40_ = 3.00, *p* = 0.047), and target step length (*F*_2,40_ = 4.51, *p* = 0.017).

### Step lengthening condition

Conditions were significantly different in target step velocity (*F*_1.56, 62.47_ = 44.07, *p* < 0.0001), trailing leg stance time (*F*_2,80_ = 21.24,*p* < 0.0001), leading leg stance time (*F*_1.35,54.11_ = 4.39,*p* = 0.027), swing time (*F*_1.59,63.78_ = 66.11, *p* < 0.0001), DS (*F*_2,80_ = 22.30, *p* < 0.0001), previous step length (*F*_1.86,74.73_ = 16.53,*p* < 0.0001), and target step length (*F*_1.44,57.67_ = 45.26,*p* < 0.0001).

Groups were significantly different in previous step velocity (*F*_2, 40_ = 5.84, *p* = 0.006), target step velocity (*F*_2, 40_ = 5.59, *p* = 0.007), trailing leg stance time (*F*_2,40_ = 59.08, *p* = 0.004), swing time (*F*_2,40_ = 3.88, *p* = 0.029), and DS time (*F*_2,40_ = 8.26, *p* = 0.001).

### Obstacle crossing condition

Conditions were significantly different in previous step velocity (*F*_1.73, 69.36_ = 17.10,*p* < 0.0001); target step velocity (*F*_1.60, 64.12_ = 6.66, *p* = 0.004); trailing leg stance (*F*_2, 80_ = 9.40, *p* < 0.0001); leading leg stance (*F*_1.68, 67.45_ = 12.22, *p* < 0.0001); swing time (*F*_2, 80_ = 39.64, *p* < 0.0001); DS time (*F*_2, 80_ = 15.014, *p* < 0.0001); target step length (*F*_1.89, 75.68_ = 42.00, *p* < 0.0001), and previous step length (*F*_1.20, 48.15_ = 8.17, *p* = 0.004).

Groups were significantly different in previous step velocity (*F*_2, 40_ = 6.27, *p* = 0.004), target step velocity (*F*_2, 40_ = 7.63, *p* = 0.002), trailing leg stance (*F*_2, 40_ = 8.77, *p* = 0.001), and DS time (*F*_2, 40_ = 7.12, *p* = 0.002). Previous step velocity in older adults with T2D was different from other groups (*p* = 0.004 and *p* = 0.009).

## Discussions

This study included three groups of participants to differentiate the effects of ageing, T2D, and the combination of T2D and ageing using novel combined paradigms of virtual stepping targets and an obstacle. For the first time, participants who reported a history of falls and had neuropathy were excluded from data analyses. The results of tests showed that older adults with T2D had normal gait in baseline when no challenge was included; however, they had impaired gait adaptability, increased step length errors, and reduced toe-obstacle clearance heights in the challenging conditions.

In line with previous research [24–28], ageing and T2D did not affect gait characteristics during walking at a preferred speed when no challenge was involved. In a study reporting gait impairments [25], experimental groups were different in characteristics such as history of falls, cognition, and the distribution of gender or walking speed [28]. Even though groups were different in these characteristics, they might not show any significant differences because locomotion was unchallenged; therefore, afferent information may not be required to update efferent copies of the locomotion in the central nerve system [29].

In adaptability test conditions, the older adults with T2D walked conservatory even when the task was to walk through without adapting sagittal foot displacement trajectories. In the walk-through conditions, participants reduced their step velocities and walked with shorter steps, longer DS and stance times, which might be interpreted as being ready to suddenly respond to a condition. The walk-through condition like other conditions in adaptability tests might increase the load of attention and as such affect gait compared with the baseline condition. Participants were ready to suddenly change their ongoing gait and find an alternative foot landing position to meet the presented tasks.

Having conservative gait patterns in the walk-through condition of adaptability tests, the older adults with T2D adapted gait parameters in the previous step in response to the step shortening condition. They reduced velocity, increased the DS time, reduced the previous step length, and then shortened their target step. However, the other groups only reduced the velocity and length of the previous step. The earlier changes in the previous step in older diabetics were more pronounced, revealing that they were more affected in the adaptability tests. Our results of perturbing step lengths (i.e., presenting SSL targets) did not confirm differences in errors between healthy older and young adults, in disagreement with previous research reports [8] in which some of the participants had experienced one or two falls a year before their participation. In this study, fall history might have increased the error of step shortening, which was an exclusion criterion in our research project. Our older participants had no history of falls and were physically fit; they walked as fast as young adults with a similar SL during overground walking.

Although older adults with T2D increased the step length of both previous and target steps in the step lengthening condition, they made the largest errors in the step lengthening condition compared with other groups. The healthy older and young groups increased the target step lengths, whereas the older diabetic group increased the swing time of the target step and increased the step lengths of the previous and target steps. To process appropriate responses, older adults with T2D reduced the velocity of the previous step and increased the stance time of the leading foot. The older with and without T2D reduced the previous step velocity when they detected the task. Healthy older adults increased the target step velocity to respond to the task; however, older adults with T2D were unable to increase the velocity of the target step as much as the healthy older adults were. Thus, they had to increase the stance time of the trailing foot in the previous step to match the toe marker with a laser beam in the target step without being able to lengthen steps accurately.

Older adults with T2D showed reduced toe-obstacle clearance height when they responded to the obstacle crossing condition. They reduced step velocities and increased the DS and stance times in response to the obstacle crossing condition. However, consistent with previous research [30], these modulations of gait parameters reduced the toe-obstacle clearance height. Using a real obstacle with other conditions made the prediction of triggering the obstacle too difficult. Participants had to cross the obstacle and could not use any other strategy such as adding a short step prior obstacle avoidance as reported as a compensatory strategy in previous research [8]. Therefore, oolder adults with T2D showed a reduced toe-obstacle clearance height. Reduced step velocities in both previous and target steps and increased DS time in older adults with T2D increased the time of responding to goal tasks compared with other groups, as reported in previous research [31]. However, they were unable to have a comparable toe-obstacle clearance height with other groups.

This study has several limitations. The choice of parameters for investigation was limited to the sagittal plane. Foot placement adjustments were not investigated in the mediolateral direction since the overground gait adaptability tests could not present targets for investigating the mediolateral foot placement adjustments. Finally, the method of sampling might limit the generalisation of the findings. Further studies are required to provide insights into the gait adaptability of older adults with T2D with participants selected more randomly across the population.

In conclusion, the study has demonstrated the interaction of ageing and T2D led to impaired gait adaptability and to more conservative gait patterns in older adults with T2D. This impaired gait adaptability may place older adults with T2D at an increased risk of falls while reacting to unexpected challenges during walking. Therefore, training programs with feedforward biofeedback [32–34] can assist this population to reinforce their feedforward internal models to control leading foot trajectories by updating the efferent copy of goal tasks and reducing the occurrence of touching obstacles during walking.

## Author Statement

All authors (SM, ST, BS, RO, and RB) contributed to the conceptualization, investigation, methodology, project administration, resources acquisition, validation, original draft writing, and final draft review and editing of the enclosed manuscript submission. SM and BS contributed to the data curation, formal analysis, and data visualization. ST, RO, and RB contributed to the funding acquisition and supervision of this study. All authors contributed to the design of the study, the interpretation of data, and the preparation of the manuscript.

## Acknowledgements

We would like to acknowledge Rhett Stephens and John Izzard for assisting with designing the setup to the specific purpose of this study. This research was conducted in the Gait Biomechanics Research Group with the support of Victoria University. The funder had no role in the study design, data collection, data analysis, interpretation of data, and drafting the manuscript.

